# Notorious Novel Avian Influenza Viruses H10N8 and H7N9 in China in 2013 Co-originated from H9N2

**DOI:** 10.1101/004390

**Authors:** Liang Chen, Feng Zhu, Chenglong Xiong, Zhijie Zhang, Lufang Jiang, Rui Li, Genming Zhao, Yue Chen, Qingwu Jiang

**Author notes:** Contributed equally.

## Abstract

In 2013, two new avian influenza viruses (AIVs) H7N9 and H10N8 emerged in China caused worldwide concerns. Previous studies have studied their originations independently; this study is the first time to investigate their co-originating characteristics. Gene segments of assorted subtype influenza A viruses, as well as H10N8 and H7N9, were collected from public database. 26 With the help of series software, small and large-scale phylogenetic trees, mean evolutionary rates, and divergence years were obtained successionally. The results demonstrated the two AIVs co-originated from H9N2, and shared a spectrum of mutations in common on many key sites related to pathogenic, tropism and epidemiological characteristics. For a long time, H9N2 viruses had been circulated in eastern and southern China; poultry was the stable and lasting maintenance reservoir. High carrying rate of AIVs H9N2 in poultry had an extremely high risk of co-infections with other influenza viruses, which increased the risk of virus reassortment. It implied that novel AIVs reassortants based on H9N2 might appear and prevail at any time in China; therefore, surveillance of H9N2 AIVs should be given a high priority.

## Introduction

In 2013, two episodes of influenza emerged in China caused worldwide concerns. A new avian influenza virus (AIV) H7N9 subtype first appeared in China on February 19, 2013. As of August 31, 2013, the virus had spread to ten provinces (Anhui, Fujian, Guangdong, Hebei, Henan, Hunan, Jiangsu, Jiangxi, Shandong, and Zhejiang) and two metropolitan cities (Shanghai and Beijing). Of 134 patients with the influenza, 45 died with a fatality rate of 33⋅58%, which is much higher compared with the fatality rate (<0⋅25%) for pandemic influenza H1N1 in 2009-2010 (Li Q et al. 2014; Chen Y et al. 2013). There were no new cases reported during the period from August 31 to October 14, 2013. On October 15, 2013, a severe human case reemerged in Shaoxin City, Zhejiang Province, and as of April 11, 2014, there reported 264 severe human cases (including 4 Cantonese confirmed in Hong Kong, one Cantonese confirmed in Malaysia, and a Hong Kong girl who had Guangdong travel history), of which there were 32 deaths (http://platform.gisaid.org/epi3/frontend#f7fab”,) Access date: April 11, 2014;(http://news.sina.com.cn/c/2014-01-28/075929365580.shtml,) Access date: February 7, 2014).

On November 30, 2013, a woman aged 73 years presented with fever and was admitted to The First Hospital of Nanchang in Jiangxi Province, China. She developed multiple organ failures and died 9 days after the onset of the disease. A novel reassortant avian influenza A H10N8 virus was isolated from the tracheal aspirate specimen obtained from the patient 7 days after the onset of illness. It is the world’s first human case infected by H10N8 subtype A influenza virus (Chen H et al. 2014). On February, 2014, another dead case caused by H10N8 avian influenza was reported in Jiangxi Provinc (http://www.morningpost.com.cn/xwzx/guonei/2014-02-14/550791.shtml,) Access date: February 14, 2014).

Sequence analyses of genomes revealed that these viruses were of avian origin, with six internal segments from avian influenza A H9N2 viruses (Chen H et al. 2014; Hu Y et al. 2013). The results done by the basic local alignment search tool (BLAST, (http://blast.ncbi.nlm.nih.gov/Blast.cgi) showed most homogenous sequences of the six internal segments associated with Asian strains isolated in between 2005 and 2013. In this study, we analyzed the phylogenetic relationship between the two novel avian influenza viruses H10N8 and H7N9, and explore their commonality in origination and evolution.

## Results

### Data collecting

A total of 4593 PB2, 4535 PB1, 4785 PA, 4688 NP, 5379 MP, and NS 5410 sequences were obtained. They all affiliated to isolates established in Asia during the period from January 1, 2004 to December 31, 2013. After convergence pre-treatment by means of MEGA6, the matrixes used for large-scale phylogenetic analysis successively contained 827 PB2, 813 PB1, 827 PA, 870 NP, 908 MP, and 890 NS.

Six additional small-scale matrixes of China novel AIVs H10N8 and H7N9 contained 77 PB2, 73 PB1, 75 PA, 79 NP, 80 MP, and 80 NS gene sequences respectively.

### Large-scale/small-scale phylogenetic trees of six interior segments

According to jModeltest, nucleotide substitution models for best maximum likelihood (ML) tree were as follows:

PB2, GTR+G (AIC=14533⋅1749),
PB1, GTR+I+G (AIC=16791⋅8743),
PA, GTR+G (AIC=32747⋅0324),
NP, GTR+I+G (AIC=13637⋅1612),
MP, GTR+I+G (AIC=7529⋅0599),
NS, GTR+I+G (AIC=7215⋅4836).

Large-scale ML phylogenetic trees were constructed according to the abovementioned substitution models by MEGA6. The results showed that influenza A viruses had a remarkable tendency of clustering according to their subtype and geographical distribution. Novel AIVs H10N8 and H7N9 shared very close homogeneity, and might originated from China AIV H9N2 circulated for long time in eastern China such as Shanghai and the provinces of Zhejiang, Jiangsu, Anhui, and Shandong.

On PB2, the donors H9N2 had emerged early in 2009 in Shanghai (KC768062DK; KC779062SW, DK or SW after the number was the host spices, the same as below, DK/duck, SW/swine, CK/chicken). By the end of 2013, their analogues were broadly distributed in Jiangsu, Zhejiang, Shandong, Hunan, Guangxi, and Hebei. Moreover, in 2013, this segment was sporadic reassorted into an AIV H5N2 in Jiangsu (KF150631CK). The phylogenetic tree also demonstrated that H7N9 could at least be divided into two sub-lineages, main lineage and Guangdong lineage.

On PB1, the donors H9N2 had circulated in Zhejiang, Shanghai, Jiangsu, Shandong, Henan, Hebei, Beijing, Hunan, Guangdong, and Guangxi since 2009.

On PA, the donor H9N2 was isolated early in 2009 in Guangxi (KF367738CK). Since then, its analogous H9N2 had sustained in Shanghai, Zhejiang, Shandong, Hebei, Hunan, Gansu, and Guangxi. Furthermore, this segment even showed a spillover to AIV H6N8 in Guangxi in 2009 (JX304767DK), and H5N2 in Hebei in 2010 (JQ041396CK).

On MP, the donors H9N2 were detected in eastern China in early 2010 (JN869533CK, Jiangsu; JF906209DK, Anhui). During the period from 2010 to 2013, they maintained a continuous epidemic by means of H9N2 in Shanghai, Zhejiang, Jiangsu, Hunan, Guangdong, and also occurred in form of H5N2 in Hebei in 2010 (JQ041412CK), so did in form of H5N1 in Hunan in 2011 (CY146711DK).

On NS, the donors H9N2 were isolated early in 2007 from multiple-regions. As of 2013, their analogues distributed in Shanghai, Fujian, Zhejiang, Jiangsu, Shandong, Henan, Hebei, Beijing, Hunan, Hubei, Jiangxi, Guangdong, Guangxi, Gansu, and Tibet, and in 2009, it occurred once in Xinjiang by the form of H5N1 subtype (CY099014 Enviroment).

**Figure 1.**
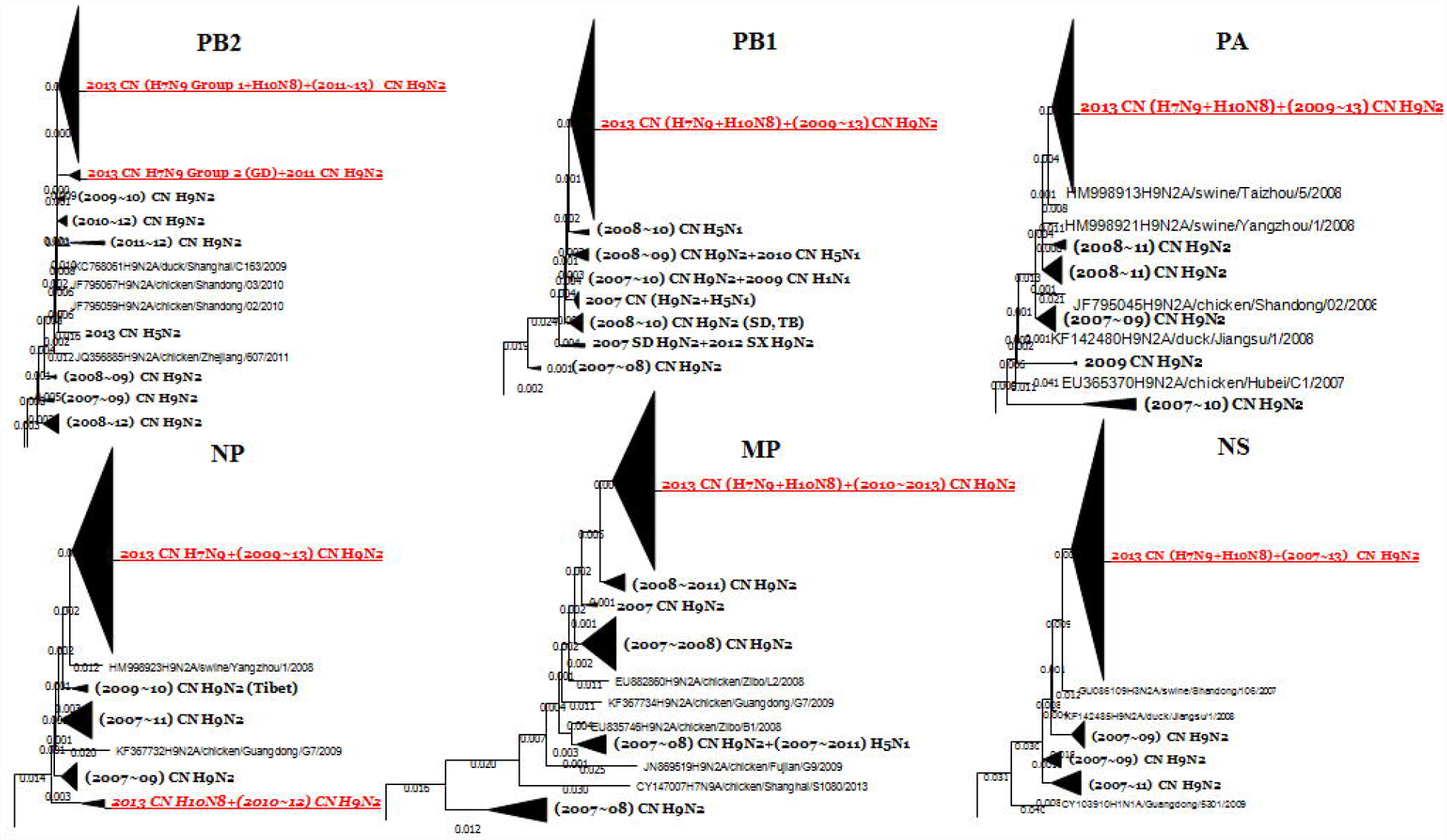
Large-scale phylogenetic trees of six interior segments affiliated to Asia influenza A viruses. All phylogenetic trees demonstrated the only branch containing AIVs H10N8or H7N9, because of too large pixels in primary plots; the latters were uploaded as supplemental figure 1. These results demonstrated that there existed very close evolutionary relationship between 2013 China novel AIVs H10N8 and H7N9, and they basically sited within the same branch on phylogenetic trees. Since these branches mainly composed of AIV H9N2 circulated in eastern and southern China without exception, novel AIVs H10N8 and H7N9 in China in 2013 were considered co-originated genetically from H9N2.

**Figure 2.**
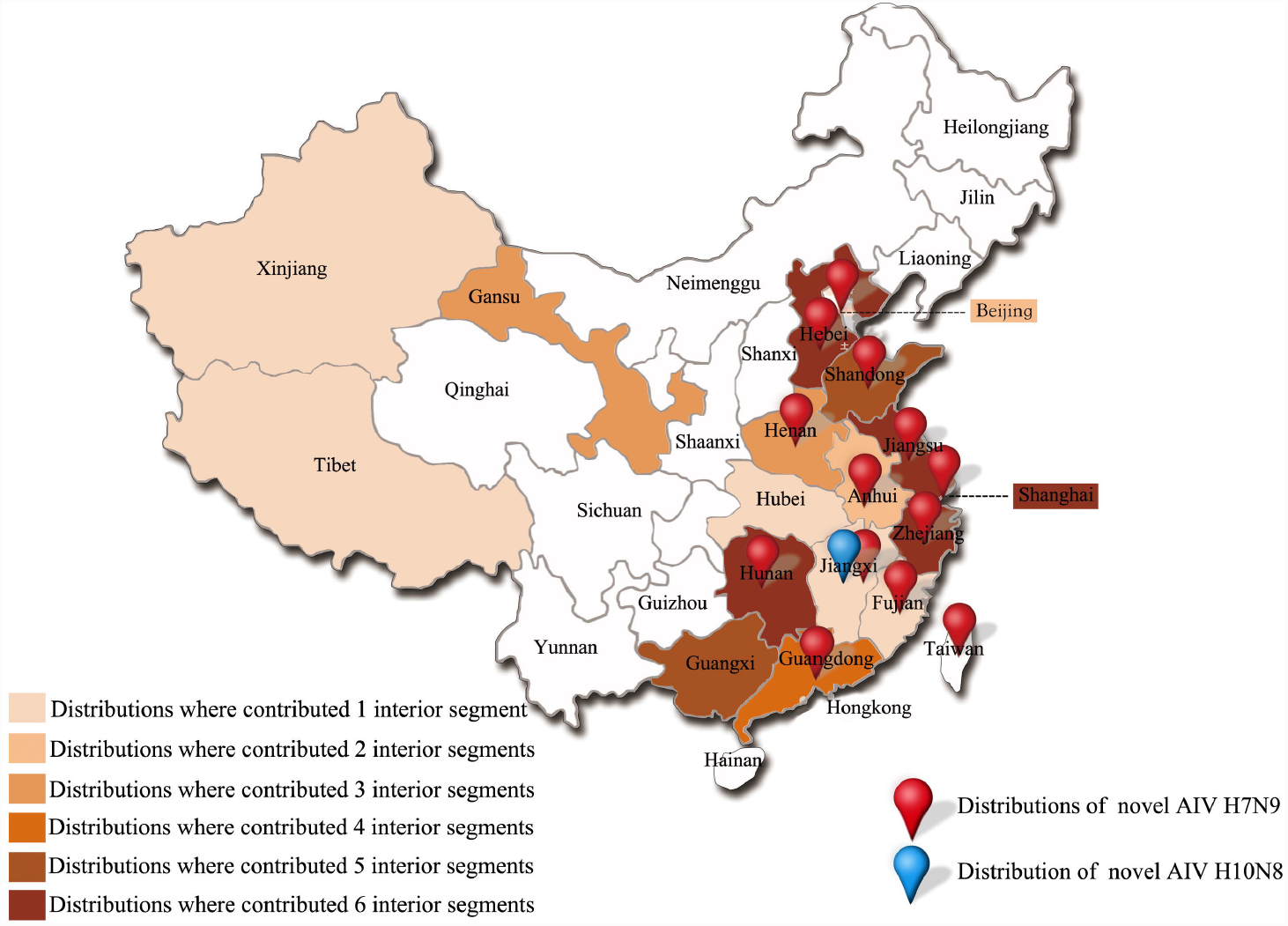
Distribution of the probable donor AIVs H9N2 and their analogues. As the probable donors for six interior segments of novel AIVs H10N8 and H7N9 in China in 2013, AIVs H9N2 and their analogues distributed extensively in eastern and southern China. Shanghai, Zhejiang, Jiangsu, Anhui, Shandong, Hebei, Hunan, and Guangdong were deeply influenced by these AIVs H9N2 and their analogues, in which, Zhejiang and Shanghai were the severest epidemic areas stricken by H7N9, whereas Jiangxi was the epidemic area of H10N8.

On NP, although it seemed that 2013 novel AIVs H10N8 and H7N9 diverged slightly from each other, they could still cluster into the same branch in the phylogenetic tree. As the donors H9N2 for H7N9, several strains were isolated in early 2009 in Shanghai (KC779054SW, KC768049DK, KC768050DK), Guangxi (KF367739CK), and Henan (KC779053SW). Zhejiang, Jiangsu, Shandong, Hunan, Guangdong, and Gansu were also their epidemic areas. The segment was also reassorted into H5N2 in Hebei in 2010 (JQ041404CK). As to H10N8, the donors H9N2 and its analogues were isolated Anhui (JF906207CK) in 2010 and Hunan (KF714784CK, 2011; KF714776CK, 2012) in 2011 and 2012 (Fig. 1 and Fig 2, Supplemental Fig. 1).

### Mean rate and tMRCA

**Table 1.** Mean evolutionary rates, tMRCA, and other parameters about six interior gene segments.

| Segment | CDS length<br>( bp ) | Mean rate<br>( 10-3/Yr.bp ) | 95% HPD<br>( 10-3/Yr.bp ) | Allowable maximum<br>variation ( bp/Yr ) | Allowable maximum<br>distance (cM/Yr) | tMRCA* | 95% HPD* |
| --- | --- | --- | --- | --- | --- | --- | --- |
| PB2 | 2280 | 4.12 | 2.85, 5.40 | 12.32 | 0.0054 | 2011.31 | 2010.68, 2011.87 |
| PB1 | 2274 | 4.01 | 3.26, 4.74 | 10.78 | 0.0047 | 2011.81 | 2011.39, 2012.23 |
| PA | 2151 | 3.36 | 2.54, 4.46 | 9.59 | 0.0045 | 2011.59 | 2011.06, 2012.09 |
| NP | 1497 | 3.16 | 2.56, 3.78 | 5.66 | 0.0038 | 2012.28 | 2011.99, 2012.54 |
| MP | 982 | 3.75 | 2.85, 4.78 | 4.69 | 0.0048 | 2012.09 | 2011.62, 2012.51 |
| NS | 838 | 2.92 | 1.93, 3.96 | 3.32 | 0.0040 | 2011.74 | 2011.16, 2012.26 |

\*Note, tMRCA of 2013 AIVs H10N8 and H7N9, and their corresponding 95% HPD values were computed based on the smaller scale matrixes only containing 2013 H10N8 and H7N9 segments, while the rest parameters were obtained based on large scale matrixes.

Six interior segments of influenza A viruses isolated in Asia on large-scale had their mean evolutionary rate ranged 2⋅92 × 10^-3^/Yr.bp ∼ 4⋅12 × 10^-3^/Yr.bp in 2004 - 2013, among which, PB2 and PB1 had a faster mutation rate compared with NS and NP (Tab. 1).

**Figure 3.**
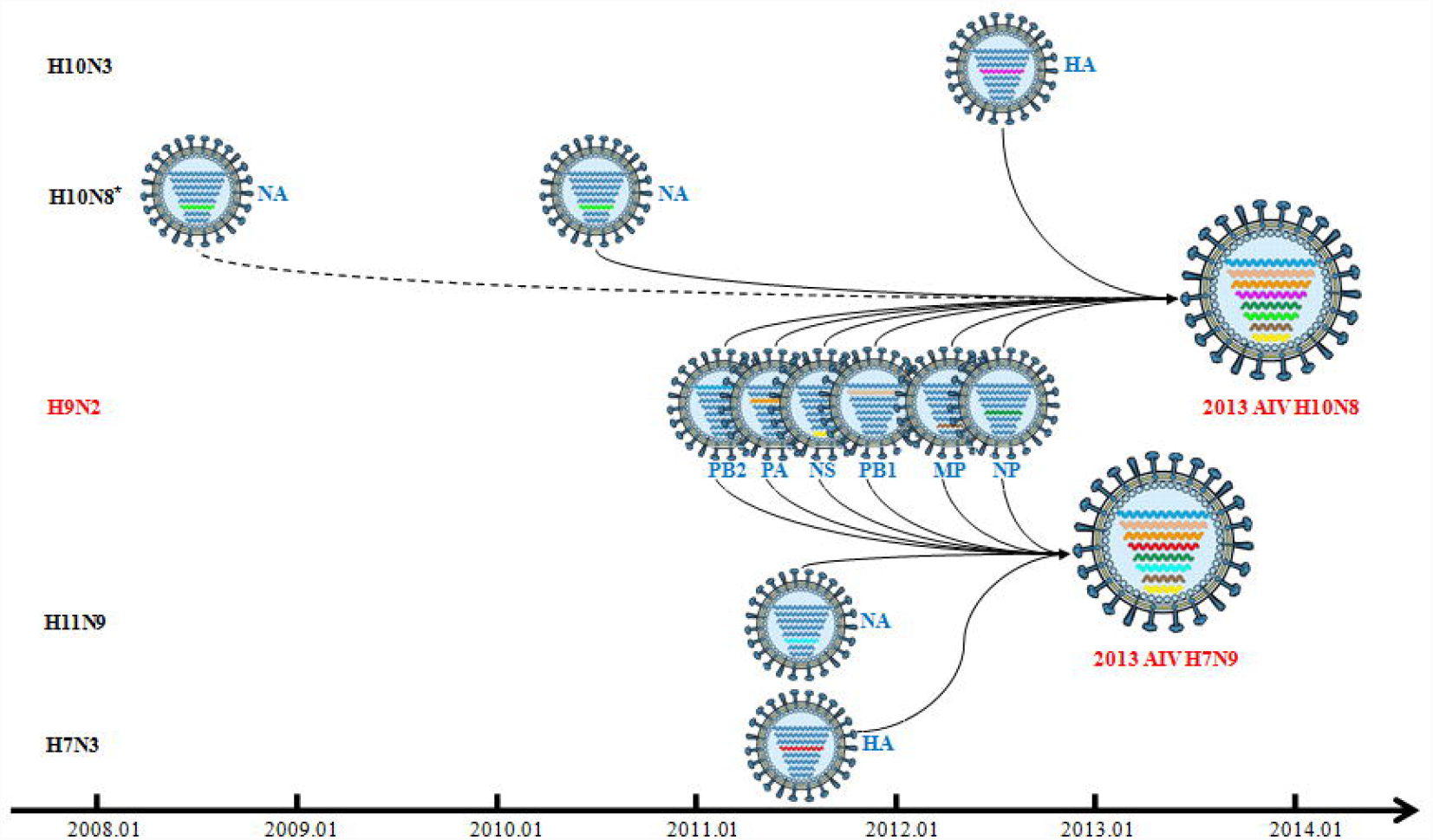
Probable evolutionary history of AIVs H10N8 and H7N9 in China in 2013. Displayed the probable procedure by this way novel AIVs H10N8 and H7N9 in China in 2013 had evolved themselves. Here just discussed the fact that the novel AIVs H10N8 and H7N9 in China in 2013 derived their six interior segments from H9N2, regardless whether or not these AIVs H9N2 were the same strain. HA and NA segments consulted references 1, 2, and 5. According to the references, HA segments of H10N8 uncertainly derived from two H10N8 isolated in different years.

According to the allowable maximum variations and allowable maximum distances, each of the six small-scale ML phylogenetic trees containing merely AIVs H10N9 and H7N9 should be divided into more than 8 clades (Supplemental Fig. 2), and these patterns must not be formed within one year. In other words, sequences in each small-scale matrix of the six interior segments must have common ancestors earlier than 2013. Analysis of tMRCA (the time to the most recent common ancestor) confirmed this inference: the most recent common ancestor of China novel AIVs H10N8 and H7N9 all emerged before March 2012, and many segments, such as PB2, might had reassorted in H9N2 in early 2011, and then into these two novel AIVs (Tab 1, Fig. 3).

### Deduced amino acid alignments

**Table 2.** Key mutations in deduced proteins coded by the six interior segments of 2013 AIVs H10N8 and H7N9.

|  |  | Known important mutation |  |  | Other sites |
| --- | --- | --- | --- | --- | --- |
|  |  | H7N9 |  | H10N8 | H10N8 differed |
|  |  | Human | AV&EV* |  | from H7N9 |
| PB2 (viral replication) |  |  |  |  |  |
| *Leu89Val* | Enhanced polymerase activity | 29 of 29 | 46 of 46 | 1 of 1 | *Asp87Glu*, |
| *Glu627Lys* | Improved viral replication | 20 of 29 | 0 of 46 | 1 of 1 | *Asp195Gly*, |
|  | at 33°C |  |  |  | *Arg340Lys*, |
|  |  |  |  |  | *Ile411Val*, |
| *Asp701Asn* | Mammalian adaptation | 2 of 29 | 0 | 0 | *Ala588Val* |
| PB1 (viral replication) |  |  |  |  |  |
| *His99Tyr* | Enables droplet transmission | 0 | 0 | 0 | *Val14Ala*, |
| *Ile368Val* | in ferrets | 23 of 26 | 42 of 45 | 1 of 1 | *Lys52Asn*, |
| PB1-F2 (induce cellular apoptosis and inhibit function of type I interferon) |  |  |  |  | *Thr291Ala*, |
| Full-length | Needed for virulence in mice | 21 of 26 | 31 of 45 | 0 | *Asn642Ser*, |
|  |  |  |  |  | *Lys757Asn* |
| Matrix protein M1 (viral assembly and budding) |  |  |  |  |  |
| *Asn30Asp*, | Increased virulence in a mice | 31 of 31 | 47 of 47 | 1 of 1 | 0 |
| *Thr215Ala* | model | 31 of 31 | 47 of 47 | 1 of 1 |  |
| Matrix protein M2 |  |  |  |  |  |
| *Ser31Asn* | Amantadine resistance | 31 of 31 | 47 of 47 | 1 of 1 | *Arg12Lys* |
| NS1 (counteracts host antiviral response) |  |  |  |  |  |
| *Pro42Ser* | Increased virulence in mice | 31 of 31 | 47 of 47 | 1 of 1 | *Ser206Gly* |
| NP (Nucleoprotein) |  |  |  |  |  |
|  |  |  |  |  | *Ile197Asn*, |
|  |  |  |  |  | *Ile217Asn*, |
|  |  |  |  |  | *Asn482Ser* |
| PA (viral replication) |  |  |  |  |  |
|  |  |  |  |  | *Ala343Ser* |

\* Note: AV=avian; EV=environment

2013 China novel AIVs H10N8 and H7N9 shared a spectrum of mutations in common on many key sites, e.g., Leu89Val and Glu627Lys on PB2; Ile368Val on PB1; Asn30Asp and Thr215Ala on M1; Ser31Asn on M2; and Pro42Ser on NS1, and so on. Especially in PB2, similar to many human derived H7N9 viruses, H10N8 had a variation of Glu627Lys. This motif, to a certain extent, was thought to be a characteristic for the human tropism of AIVs, and could improve viral replication at 33°C. On some other sites, deduced amino acid of H10N8 differed from that of H7N9 (Tab. 2).

## Discussion

The novel AIVs H10N8 and H7N9 identified in China in 2013 had a common origin. AIV H9N2 circulated everlastingly in eastern China had donated six interior segments for their genomes. Their interior segments had evolutionarily achieved in H9N2 likely in 2011-2012. Having accomplished two spillovers from poultry to human being by means of H10N8 and H7N9 in such a short period, H9N2 demonstrated its high frequency and efficiency on virus reassortment.

The two notorious novel AIVs shared a spectrum of mutations in common on many key sites implied that they might have the similar pathogenic, tropism and epidemiological characteristics. It is important to strengthen the surveillance and to reinforce case management of novel AIV H10N8 and novel AIV H7N9.

The first strain of H9N2 AIV was isolated in 1966, and soon a large number of infections caused by H9N2 in poultry were reported (Homme P and Easterday B.1970). In 1988, Perez et al (2003) established three H9N2 AIVs from quail in Hong Kong, this is the first time of confirming H9N2 prevalence in Asia poultry. In mainland China, the emergence of H9N2 AIV was appeared in 1994 (Guo X et al. 2003). So far in southern China and Hong Kong, H9N2 are divided into three sub-lineages, A/Chicken/Hong Kong/G9/97(H9N2) (G9-Like), A/Quail/HongKong/G1/97(H9N2) (G1-Like), and A/Duck/HongKong/Y439/97(H9N2) (Y439-Like). Further monitoring showed that two lineages of H9N2 influenza viruses, A/Quail/Hong Kong/G1/97 (Qa/HK/G1/97)-like and A/Duck/Hong Kong/Y280/97 (Dk/HK/Y280/97)-like, widely distributed in the live-poultry in southeastern China. More than 16% of cages of quail in the poultry markets contained Qa/HK/G1/97-like viruses, and the Qa/HK/G1/97-like viruses were evolving rapidly, especially in their PB2, HA, NP, and NA genes (Guan Y et al. 2000). In this study, H9N2, as the donor for 2013 China novel AIVs H10N8 and H7N9, is genetically Qa/HK/G1/97-like.

H9N2 acting as a donor and providing gene segments to reform a novel subtype influenza virus is a very common phenomenon. Gu et al (2010) isolated one H5N1 A/duck/Shandong/009/2008 from an apparently healthy domestic ducks in eastern China in 2008. According to their BLAST results, four interior gene segments (PB2, PB1, PA and M) of this isolate displayed the closest relationship with H9N2 subtype that was prevalent in eastern China, and then this H5N1 was thought as a reassortant virus derived from G1-like H9N2 and H5N1 subtypes. Zhang et al (2009) studied the genetic and antigenic characteristics of H9N2 influenza viruses isolated from poultry in eastern China during the period from 1998 to 2008 and suggested that the Ck/SH/F/98-like H9N2 virus may have been the donor of internal genes of human and poultry H5N1 influenza viruses circulating in Eurasia. Experimental studies showed that some of these H9N2 viruses could be efficiently transmitted by the respiratory tract in chicken flocks. Gene exchanging, i.e. H9N2 derives their segment from other subtype influenza viruses, also was found. Cong et al (2007) reported five swine H9N2 influenza viruses isolated from diseased pigs from different farms possessed H5N1-like sequences of the six interior genes, indicating that they were reassortants of H9 and H5 viruses. Reassortant H9N2 influenza viruses containing H5N1-like PB1 genes in southern China were reported by Guan et al (2000) and Dong et al (2011). Such a high frequency of gene exchanging between H9N2 and other subtype influenza viruses implied that novel AIV reassortants based on H9N2 might appear and prevail at any time; therefore, surveillance of H9N2 AIVs, specially, H9N2 AIVs in eastern China, should be given a high priority.

Reassortment contributing to the generation of genetic diversity can only occur among viruses, which replicate within same cells. The prerequisite for reassortment is that an individual host simultaneously infected with multiple divergent viral strains and then formed a quasispecies pool consisting of rather closely related members (Padidam M et al. 1999; Robertson D et al. 1995). For a long time, three genetic sub-lineages of H9 virus (G1, G9, and Y439) had been circulated in eastern and southern China, and poultry was the stable and lasting maintenance reservoir (Peiris M et al. 1999; Guo Y et al. 2000). The perennial positive rate of antibody against H9N2, as a typical low pathogenic avian influenza virus and a major contributor to this novel H7N9, fluctuates between 5•3% and 12•8%, and the rate of virus isolation could even reach 9% in poultry, but this did not cause any obvious epidemic with mass poultry deaths (Cheng X et al. 2002; Lin Y et al. 2002). On April 3, 2013, during the epidemic of H7N9, we collected the last samples from live poultry market in Shanghai (on April 4th, Shanghai closed the live bird market); from 300 anal swabs of chicken, goose, duck and pigeon, we detected 7% influenza A virus positive, and all of them were H9N2 subtype. On December 25, 2013, in Jiangsu, another eastern China province, from 268 anal swabs of chicken, goose, duck and pigeon, and fecal samples of wild bird, we obtained 14 H9N2 positives, no other subtype (unpublished). Such a high carrying rate of AIV H9N2 in poultry had an extremely high risk of co-infections with other influenza viruses, which increased the risk of virus reassortment. In fact, the reassortment between H5N1 and H9N2 had frequently occurred in China as abovementioned. Our study hereby provided evidence for the six interior segments had appeared in 2012 or before, and later reassorted into the novel H10N8 and H7N9 AIVs, which caused an epidemic in 2013. So, it is important to strengthen the surveillance and to reinforce case management of novel AIV H10N8 and novel AIV H7N9.

H9N2 AIVs act not just donors for assembling a novel influenza virus and they could act also as pathogens to contaminate mammalians directly. H9N2 is the only one of H9 subtype could function that way (Lin Y et al. 2002; Peiris J et al. 2001). In 1998, swine H9N2 virus was isolated in Hong Kong; this is the first time of H9 subtype AIV infecting mammalian (Saito T et al. 2001). Peiris et al had investigated live pigs traded from southern China to Hong Kong in 1998-2000 and confirmed that H9N2 had been widely spread among pigs (Peiris J et al. 2001). As for H9N2, adaptation to pig was an important milestone on the way to human infection. In 1998, human H9N2 infections emerged in Shaoguan and Shantou Cities of Guangdong Province (Huang P et al. 2001). Peiris also reported two human cases in Hong Kong in 1999 (Peiris M et al. 1999). Therefore, from the point of view that H9N2 could act as potential human pathogen, there is a need for a strict monitoring of H9N2 AIVs in China.

Overall, there is an important public health risk for avian influenza A H7N9. The novel avian influenza A H10N8 might be a new wave in the future. As a typical moderate virulence AIV, H9N2 had a very high carrying rate in poultry; it would inevitably lead to a high frequency of reassortment. We deeply concerned here that the next wave of avian flu in the future is extremely likely to be caused by the reassortants of H9N2, or the variant ones from it. Influenza epidemiological and virological surveillance should be further strengthened, especially on H9N2.

## Methods

### Data collecting

Genomes of novel AIV H10N8, named as Jiangxi Donghu, were downloaded from the Global Initiative on Sharing Avian Influenza Data (GISAID) database (http://platform.gisaid.org/epi3/frontend) on February 6, 2014, with their GISAID Numbers being EPI_ISL_152846_497477 ∼ 497484. The six interior gene sequences of Asian influenza A viruses identified during 2004 to 2013 were collected from the NCBI Influenza Virus Sequence Database(http://www.ncbi.nlm.nih.gov/genomes/FLU/aboutdatabase.html), and the search formula “Type+Host+Country/Region+Protein+Subtype+Sequence length+Collection date” was set as “A+any+Aisa+PB1 ∼ NS+any+Full-length plus+2004-01-01 ∼ 2013-12-31”. Additional matrixes used for calculating the most recent common ancestor (tMRCA) of H10N8 and H7N9 were searched by the formula “A+any+China+PB1 ∼ NS+H7N9+Full-length plus+2013-03-14 ∼ 2014-02-07”. In order to facilitate the following analyses, sequences in each matrix were named as “Access No.+Host+Region/Country+Subtype+Year”.

MEGA6 (http://megasoftware.net) was used for the pre-treatment of genome data. According to the amount of gene sequences, all large-scale matrixes were split into 15 ∼ 18 sub-matrixes simultaneously containing H10N8 and representative H7N9 strain named A/Zhejiang/DTID-ZJU01/2013(H7N9). After W method alignment, a phylogenetic tree was constructed for each sub-matrix by p-distance. The sequences were kept only one or more for further analysis if 1) they are the sequences with the same HA and NA subtypes, and were isolated from the same area within 2 years, and 2)they showed high homogeneity in the phylogenetic tree, since the other might be the same strain but established from different individuals.

### Large-scale/small-scale phylogenetic trees of six interior segments

jModeltest 2.1.3 program (http://darwin.uvigo.es) was applied to estimate the model’s likelihood value and to search the best ML tree. Three substitution schemes and then 24 candidate models were evaluated.

After aligned by clustalW using the MEGA6 software, ML trees of six large-scale gene sequence matrixes were constructed, with the nucleotide substitution model selecting conducted by Akaike information criterion (AIC) values obtained from jModeltest 2.1.3. Significance testing was bootstrapped with 1000 replicates. Six small-scale ML phylogenetic trees only containing the corresponding segments affiliated to 2013 China novel AIVs H10N8 and H7N9 were constructed by nucleotide substitution model GTR +I +G; The significance testing was bootstrapped with 1000 replicates, too.

### Computation of mean evolutionary rate and tMRCA

Both mean evolutionary rate and tMRCA were calculated by using the BEAST v1.6.1(http://beast.bio.ed.ac.uk). Mean evolutionary rate was computed based on six large-scale gene sequence matrixes, while the tMRCA between AIVs H10N8 and H7N9 was deservedly deduced based on six small matrixes that only comprised of the segments affiliated to the novel H10N8 and H7N9 identified in 2013. For tMRCA analysis, the collection dates of AIVs H10N8 and H7N9 were calculated by using the following formula:

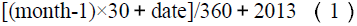

In large-scale analysis, models derived from jModeltest 2.1.3 were used again for choosing nucleotide substitution, and uncorrelated lognormal relaxed clock model was engaged as clock model, while in tMRCA analysis the model of nucleotide substitution and clock were all set as GTR + I + G and strict respectively. Log files were imported into Tracerv1.5 (http://beast.bio.ed.ac.uk) to read the needed data.

The allowable maximum variations and allowable maximum distances of each segment were deduced by the obtained data based on large-scale matrixes.

### Alignments of deduced amino acid

To compare the mutations occurred in some key sites, open reading frames (ORF) of interior segments affiliated to China novel AIVs H10N8 and H7N9 were translated into amino acid, and alignments were done by using the MEGA6.

## Data access

All raw and processed data from this study have been derived from 1), Global Initiative on Sharing Avian Influenza Data (GISAID) database (http://platform.gisaid.org/epi3/frontend); 2), NCBI Influenza Virus Sequence Database (http://www.ncbi.nlm.nih.gov/genomes/FLU/aboutdatabase.html).

## Acknowledgments

We thank all the scientists for having subscribed the genomes of AIVs H10N8 and H7N9 to NCBI Influenza Virus Sequence Database and the Global Initiative on Sharing Avian Influenza Data (GISAID) database.

This research was supported by grants awarded to Prof. Jiang from National Natural Science Foundation of China (grants No. 81172609), Prof Zhao from the Key Discipline Construction of Public Health of Shanghai of Shanghai Municipal Health and Family Planning Commission (grants No. 12GWZX0101). It also supported by Dr. Zhijie Zhang’s grants of the Ecological Environment and Humanities/Social Sciences Interdisciplinary Research Project of Tyndall Center of Fudan University (FTC98503A09). The funders had no role in the study design, data collection and analysis, the decision to publish, or the preparation of the manuscript.

## Disclosure declaration

The authors declare they have no competing interests.

## Figure legends

**Supplemental figure 1.** Whole plot large-scale ML phylogenetic trees of six interior segments affiliated to Asia influenza A viruses. Corresponding to Figure 1, novel AIVs H10N8 and H7N9 in China in 2013 co-originated genetically from H9N2 circulated in eastern China. Branches pointed by arrow were those ones on which the 2013 China novel AIVs H10N8 or/and H7N9 sited.

**Supplemental figure 2.** ML phylogenetic trees of six interior segments affiliated to 2013 AIVs H10N8 and H7N9. According to the allowable maximum variations and allowable maximum distances, each tree of six interior segments affiliated to 2013 AIVs H10N8 and H7N9 should be divided into more than 8 clades.

